# A history of symbiosis impacts the host evolutionary trajectory in experimentally evolved amoebas

**DOI:** 10.64898/2026.03.18.712585

**Authors:** Israt Jahan, Tyler J. Larsen, Joan E. Strassmann, David C. Queller

**Author notes:** Corresponding author and email address; Israt Jahan. The authors confirm all supporting data, code, and protocols have been provided within the article or through supplementary data files.

## Abstract

Biological diversity driven by endosymbiosis arises from the intertwined evolution of microbes and their hosts. Each partner affects the fitness and therefore the evolution of the other. Here, we tested a further question: does the history of symbiosis affect evolution even after the partnership is dissolved? We analyzed phenotypic data from experimentally evolved strains of *Dictyostelium discoideum* hosts, each of which had had its symbiont removed, to study how their traits evolved. We found that host trait evolution was affected by the prior history of infection, specifically by which of three *Paraburkolderia* bacterial symbionts had been removed. Thus, symbionts affect not only current evolution but also generate path dependence that affects the subsequent evolutionary trajectories even after the symbionts are lost.

**Impact statement:** The evolution of partner dependence in host-microbial symbioses has fundamentally shaped biological diversity and ecosystem function. To examine variation in symbiont dependence in the social amoeba, we compared how different strains of *Dictyostelium discoideum* respond evolutionarily after the loss of their bacterial symbionts. We analyzed phenotypic data from experimentally evolved strains and found that the absence of different symbiont species leads to distinct changes in the subsequent evolution of key traits like cell proliferation, slug migration, and spore production. This research expands our current understanding of microbial symbiosis by revealing that symbiont species may impact the evolution of their hosts even after the symbiont is gone.

**Data summary:** We used phenotypic traits data from our previous experimental-evolution dataset from the open-access repository Dryad (https://doi.org/10.5061/dryad.kkwh70s97). Scripts for the statistical analyses are available in a GitHub repository (https://github.com/jahanisrat/SymbiontLoss). The accompanying R project includes code to reproduce the graphs in the results section.

## Introduction

Symbiosis is a strong molder of both the ecology and evolution of life on earth (1–5). It can open up new ecological opportunities for the participants, as mycorrhizae did for land plants (6–9). Loss of symbionts can also trigger cascading effects across ecological scales, as demonstrated by the effects of coral bleaching where environmental stress disrupts partnerships between corals and their photosynthetic algae (10, 11). On the evolutionary side, the presence of symbiosis clearly affects fitness, typically positively for mutualistic partners but often negatively for antagonistic symbioses. This can strongly affect the course of subsequent evolution. However, the evolutionary consequences of symbiont loss remain less understood. The evolutionary history of symbiosis may impose historical contingencies on the host, influencing its evolutionary trajectories even after symbiont loss.

One microbial system that has emerged as a model for studying host-symbiont interdependence and its evolutionary implications is the social amoeba *Dictyostelium discoideum* and its *Paraburkholderia* endosymbionts. This soil-dwelling protist is a generalist predator (12) that participates facultatively in symbiotic associations with several species of bacteria (13–15). Its unique life cycle, which includes both unicellular and multicellular stages, adds a layer of complexity to its relationship with bacterial endosymbionts from the genus *Paraburkholderia* (16, 17). Unlike ancient endosymbioses where transitional states are obscured by time, the *Dictyostelium-Paraburkholderia* symbioses show various stages of symbiotic integration (17–19), from facultative associations to emerging dependencies (20–22).

Three novel bacterial species of *Paraburkholderia* – *P. agricolaris*, *P. hayleyella*, and *P. bonniea –* have been isolated from wild collected *D. discoideum* (20, 23, 24). The outcomes of the symbiotic relationships between *Paraburkholderia* and its host *D. discoideum* range from mutualism to parasitism, sometimes by the same symbiont species (25, 26). Two of the endosymbionts – *P. bonniea* and *P. hayleyella* – are sister species that have highly reduced genomes compared to other members of the genus, including the other symbiont, *P. agricolaris.* These two reduced-genome bacteria have lost much, but not all, of their metabolic independence, relying on their host for essential nutrients and exhibiting limited carbon metabolism (13). Taken together, it is likely that the reduced symbionts have a long-term co-evolutionary relationship with their host. In contrast, *P. agricolaris* has retained a more complete genome and is closely related to species that are not symbiotic with *Dictyostelium*, suggesting a less intimate or shorter-term association with its host and greater ecological flexibility.

Experimental evolution approaches have emerged as powerful tools for examining the selective pressures and constraints that shape symbiotic relationships over time (27, 28). A previous study by Larsen et al. combined experimental evolution and disruption of established symbioses to reveal adaptations that help maintain complex social interactions in the wild (29). In one analysis, which did not involve the symbionts, the authors showed that social traits in wild-collected *D. discoideum* strains tend to deteriorate when evolved in unicellular non-social conditions in the laboratory (29) suggesting that those traits were adaptations. A similar examination of trait deterioration in *Paraburkholderia* in the absence of their hosts suggested that some traits were adaptations that helped or harmed the host, and others showed no evidence for either (30).

Here we investigated a different question using the same experimentally evolved strains: how does symbiotic history shape subsequent host evolution in the social amoeba after loss of the symbiont? By analyzing phenotypic data from the laboratory-evolved *D. discoideum* lineages (29, 31), we tested whether having previously maintained a symbiont influenced the magnitude and direction of subsequent evolutionary responses in three key *D. discoideum* life-history traits: cell proliferation, migration distance, and spore production. We examined host strains that had evolved after being cured of their symbiont species to those that did not carry symbionts to begin with. We further compared the overall evolutionary response of the strains to test whether specific symbiont species that *D. discoideum* previously harbored – *P. agricolaris*, *P. hayleyella*, or *P. bonniea –*shaped distinct aspects of host evolution. Thus, we ask whether symbiotic associations can result in historically contingent effects on the evolutionary trajectory of host *D. discoideum* even after the symbionts are removed.

## Results

### Prior history of symbiosis does not have general effects on subsequent evolution in *D. discoideum*

We asked whether all *D. discoideum* strains that had been cured of their *Paraburkholderia* symbionts (called symbiotic strains) evolved differently from *D. discoideum* strains that did not carry any symbionts prior to experimental evolution (non-symbiotic strains). There are multiple ways of examining this question. First evaluate each trait individually, asking both whether strains that had lost their symbionts, regardless of species, evolved differently from non-symbiotic strains, and then whether the strains behaved differently according to which symbiont species they had lost. Finally, we examine the multi-trait responses.

To understand individual trait evolution and its interaction with the history of symbiosis, we generated linear mixed effect models individually for the three traits – cell proliferation, migration distance, and spore production (model M1, Methods: Statistical analysis). First, we investigated which traits showed evidence of evolutionary change during the experiment. Estimated marginal means revealed that in hosts with a history of symbiosis, evolved lineages showed higher cell proliferation, higher spore production, and shorter migration distances than their ancestors over the evolutionary period without their symbionts. In contrast, in hosts without a symbiotic history, there was no significant difference between evolved and ancestral lines (Table 1). This seems to suggest that past association with a *Paraburkholderia* symbiont, regardless of the species harbored, affects subsequent evolution after the symbiont is lost.

**Table 1.** Statistics from pairwise contrasts (t-tests, adjusted for multiple comparisons using FDR method) between Ancestors and Evolved Lineages for univariate *Dictyostelium discoideum* traits. The traits cell proliferation and spore production were calculated as log transformed number of cells and spores respectively. Slug migration was measured as distance traveled towards light in centimeters.

| Trait | Symbiotic Status | Estimated contrast<br>(Evolved – Ancestor) | SE | z.ratio | p.value |
| --- | --- | --- | --- | --- | --- |
| Cell Proliferation | Non-symbiotic | 0.073 | 0.083 | 0.878 | 0.380 |
| **Cell Proliferation** | **Symbiotic** | 0.118 | 0.054 | 2.166 | **0.030** |
| Slug Migration | Non-symbiotic | -0.143 | 0.506 | -0.282 | 0.778 |
| **Slug Migration** | **Symbiotic** | -0.687 | 0.331 | -2.073 | **0.038** |
| Spore Production | Non-symbiotic | 0.041 | 0.078 | 0.522 | 0.602 |
| **Spore Production** | **Symbiotic** | 0.105 | 0.051 | 2.061 | **0.039** |

However, though none of the non-symbiotic traits changed significantly, they all moved to some extent in the same direction as in the symbiotic lineages. The changes from ancestor to evolved in the two groups can therefore be better compared with a difference-of-differences contrast from the estimated marginal means. The difference in evolutionary response between symbiotic and non-symbiotic hosts was not significant in any of the traits (Supplementary Table S1), indicating that although evolution affected symbiotic hosts, the magnitude or direction of this effect was not significantly stronger than in non-symbiotic hosts. This conclusion was also supported by the non-significant interaction between treatment and symbiosis in all three models (Supplementary Table S2A).

### Prior history of particular symbionts affects subsequent evolution of cell proliferation in *D. discoideum*

The analysis above comparing non-symbiotic strains to symbiotic strains pooled together the strains that lost three species of symbionts (Model M1, Fixed effects: Treatment*Symbiosis). This may have masked some underlying variation due to the species of symbiont. To investigate whether changes in specific traits differed according to which symbiont species they lost, we fitted another robust linear mixed model for each trait taking the lost symbiont species into account (M2, Methods: Statistical analysis, Fixed effects: Treatment*Symbiont). In this model, the interaction term is significant for cell proliferation (Supplementary Table S2B). This indicates different evolutionary responses among the four symbiont treatments, including the natively symbiont-free treatment. that lacked symbionts from the beginning. The interactions for the other two traits were not significant. Interaction plots are shown in Figure 1A.

**Figure 1:**
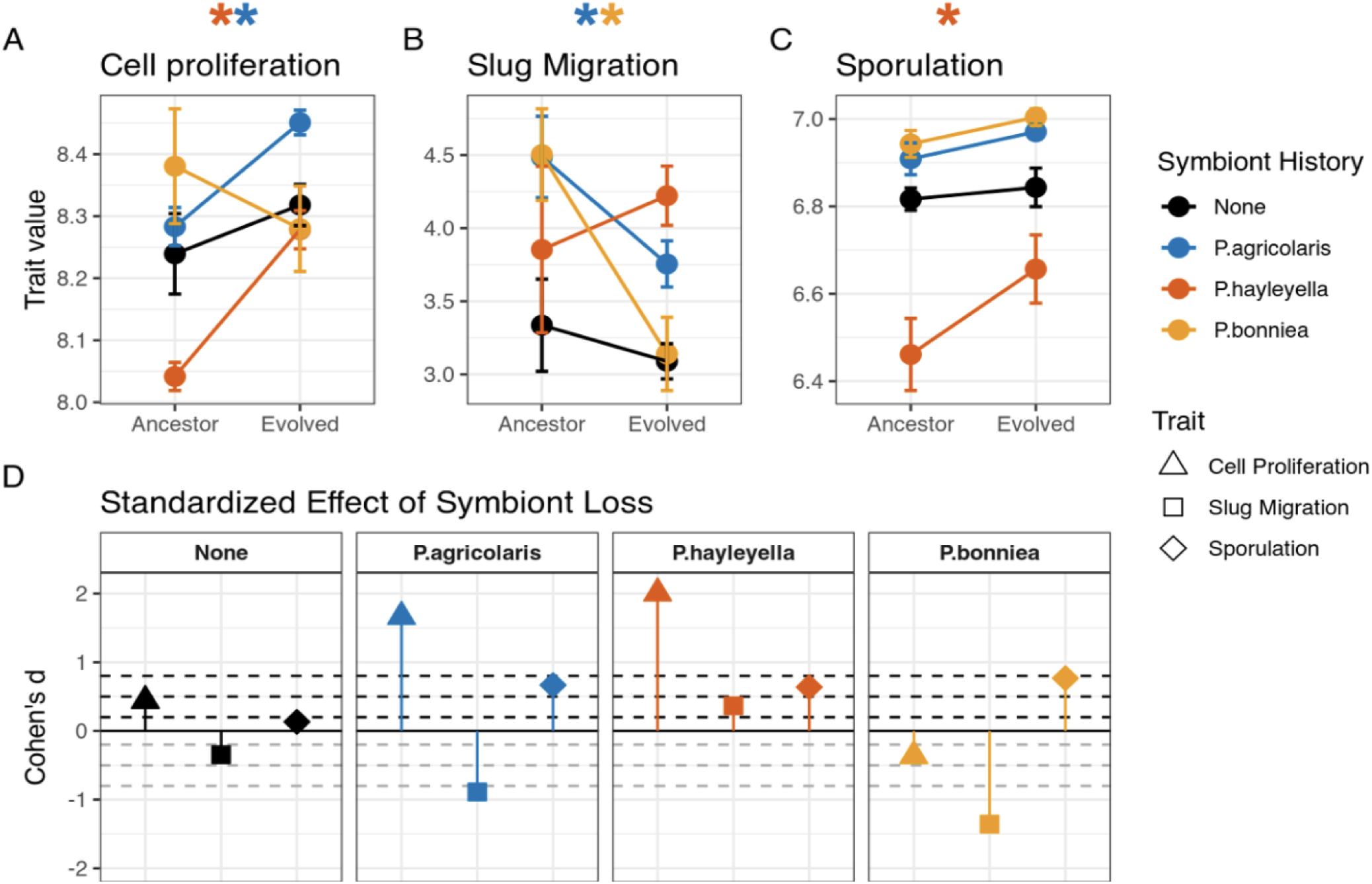
*Dictyostelium discoideum* strains that host symbionts in the wild show distinct evolutionary responses after the loss of symbiosis. Top panels (A-C) show average phenotypic means and their standard errors for Ancestor and Evolved strains. The three traits in this study are (A) Cell Proliferation, (B) Slug Migration (C) Spore production. The asterisks correspond to the traits that show a significant change after experimental evolution. The bottom panels (D) show standardized effect sizes (Cohen’s d) for each trait (cell proliferation, slug migration, spore production) by symbiont history. The dashed lines represent the threshold for effect sizes (small: 0.2, intermediate: 0.5, large: 0.8). Lines above zero indicate a positive effect on trait values following symbiont loss, while lines below zero represent a negative effect.

We examined loss of individual symbiont species using estimated marginal means and post-hoc pairwise contrasts (see Methods). For cell proliferation, evolved lines exhibited higher proliferation than their ancestors for lines in the absence of the symbiotic partner *P. agricolaris* (Evolved − Ancestor = 0.173 ± 0.077, z = 2.267, p = 0.023) and *P. hayleyella* (Evolved − Ancestor = 0.237 ± 0.094, z = 2.528, p = 0.011), as visualized in Figure 1A. We observed a decline in slug migration with evolution in lines that evolved without *P. bonniea* (Evolved − Ancestor = −1.496 ± 0.648, z = −2.308, p = 0.021; Figure 1B). Spore production significantly increased in lines that evolved without *P. hayleyella* compared to their ancestors (Evolved − Ancestor = 0.205 ± 0.094, z = 2.171, p = 0.03; Figure 1C). We present a systematic account of the pairwise contrasts within strains with a history of carrying a specific symbiont in the Supplementary Table S3.

We further standardized trait values using effect sizes (Cohen’s d) to compare the response to experimental evolution across traits and symbiont histories. The non-symbiotic strains showed negligible changes in any trait (Cohen’s |d| < 0.2) while strains that had been cured of their natural symbionts showed a clear evolutionary response in at least one trait per symbiont species (Figure 1D). These standardized effect sizes reveal that even when individual traits show only modest statistical significance, several respond consistently in the same direction under experimental evolution.

### Symbiont history affects multivariate evolution of host traits and evolutionary tradeoffs

To test whether strains with a history of symbiosis differed significantly in their overall trait responses, we performed multivariate analyses on all three traits combined (cell proliferation, slug migration, spore production). First, we performed a Principal Component Analysis (PCA) to understand the magnitude and directionality of evolutionary change in trait space. The first two principal components captured approximately 71% of the total variation in *D. discoideum* traits (PC1: 36.44%, PC2: 34.73%). PC1 was strongly negatively associated with cell proliferation (−0.767), moderately negatively associated with slug migration (−0.527), and weakly negatively associated with spore production (−0.366). PC2 captured a trade-off between spore production and slug migration distance, with strong negative loading on spore production (−0.768) and strong positive loading on slug migration (0.636), while cell proliferation had minimal influence (−0.071). This axis appears to represent a resource allocation trade-off between reproductive output and dispersal capability. Strains with different symbiont histories occupied distinct regions of this trait space (Figure 2A), with evolved populations showing consistent directional shifts relative to their ancestors.

**Figure 2:**
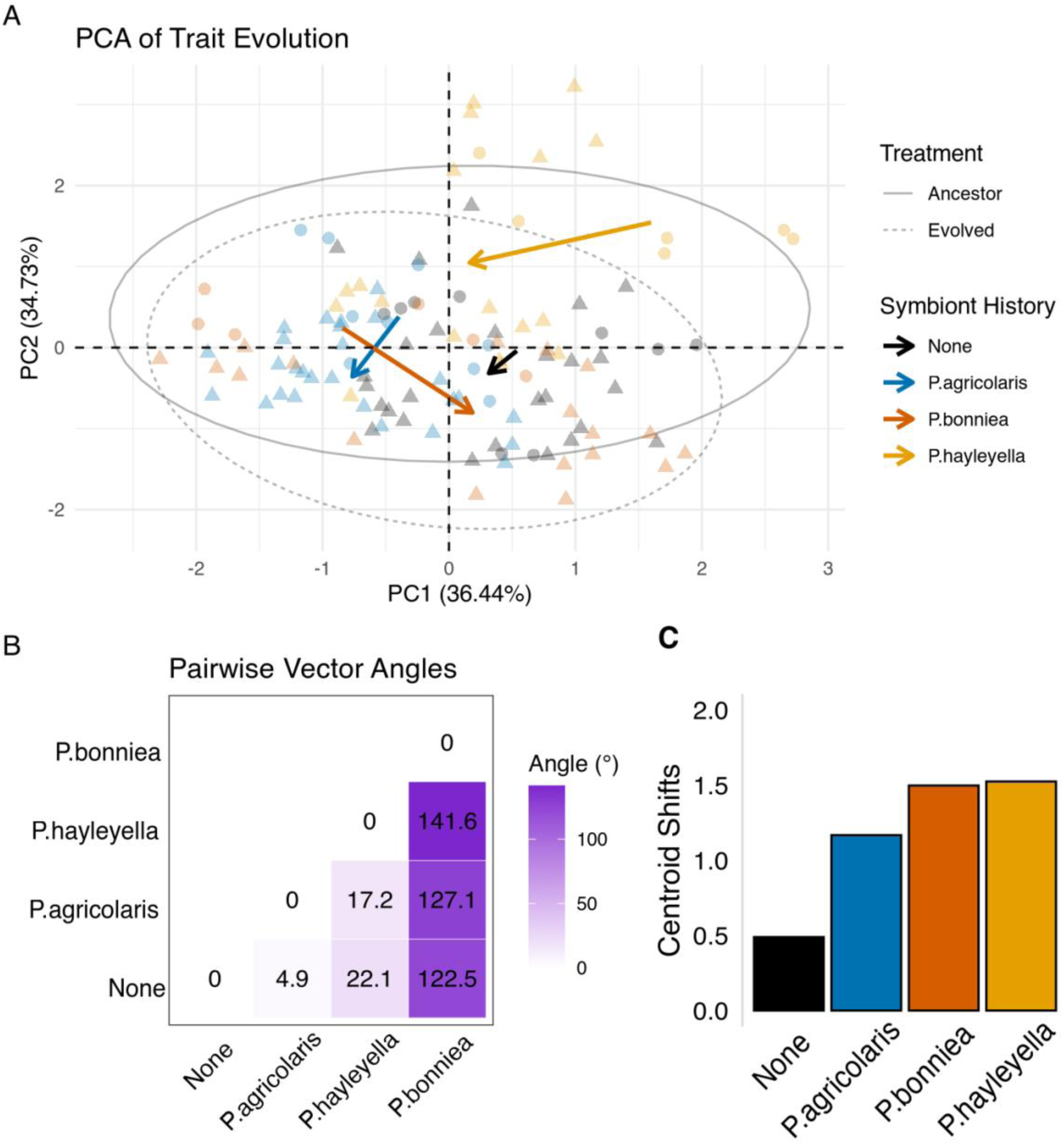
Panels show evolutionary trajectories for each symbiont history (None, *P. agricolaris*, *P. bonniea*, and *P. hayleyella*). (A) PCA scores for each data point. Ellipses encompass the spread of data (95% confidence), and arrows indicate the direction and magnitude of evolutionary change between the centroids of ancestral and evolved treatments. Circles represent ancestral populations, and the triangles represent evolved populations. Points are colored by the symbiont the strains associated with in the wild, (B) Heatmap shows pairwise angular deviations between the strain types in multivariate trait space. Angular deviation was calculated from vectors representing shift in traits from ancestors to evolved for each strain type. (C) Bar plots showing the magnitude of evolutionary change in trait space (all three PCs were included).

Next, we examined shift in position along all three principal components to identify patterns of adaptation associated with different symbiont histories. The evolutionary trajectories, represented by vectors from ancestral to evolved centroids revealed that symbiont history not only influenced the magnitude of change but also its direction in multivariate trait space (Figure 2A-C). We quantified symbiont-specific evolutionary shifts by comparing angular deviations (Figure 2B) and centroid shifts along all three principal components on the trait space (Figure 2C). The angles and vector magnitudes highlight that hosts that lost *P. bonniea* and *P. hayleyella* exhibit the most deviation from non-symbiotic strains. Hosts that lost *P. agricolaris* evolved in rather similar manner to the non-symbiotic hosts. *P. bonniea* hosts and *P. hayleyella* hosts showed the largest centroid shifts. Despite losing symbionts that are sister species, these hosts evolved in very different directions from each other.

We then performed a two-way MANOVA (Multivariate Analysis of Variance) with Treatment (Ancestor or Evolved), symbiont history and their interaction as predictors to test for overall significance of effects of symbiont loss. The multivariate analysis revealed significant effects of experimental evolution (treatment) (Pillai’s trace = 0.180, p < 0.001), symbiont history (Pillai’s trace = 0.668, p < 0.001), and their interaction on the combined traits (Pillai’s Trace = 0.172, p = 0.018). In addition to the standard F-distribution-based tests, we calculated Chi-square approximations for our multivariate test statistics to ensure robust interpretation (Supplementary Table S5).

We further confirmed these findings using a PERMANOVA analysis with the same predictors with strain-level stratification. We observed significant effects for treatment (p < 0.001) and symbiont history (p < 0.001). Most importantly, the treatment × symbiont interaction remained significant (p < 0.05) even after controlling for strain-specific variation. This robust statistical approach accounts for non-independence of observations from the same strains and reinforces our conclusion that the evolutionary response to symbiont-free conditions significantly depends on previous symbiont associations.

## Discussion

The concept of historical contingency suggests that past interactions and events influence subsequent evolutionary trajectories and possibilities through the accumulation of path-dependent changes (27, 28, 32). The role of historical contingency is particularly relevant to the evolution of symbiosis (33, 34). Recent theoretical work suggests that historical symbiotic associations may create evolutionary “lock-in” effects, where initial facultative partnerships can lead to obligate dependencies through the gradual accumulation of complementary traits and loss of redundant functions (35, 36).

Previous studies on the *Dictyostelium-Paraburkholderia* symbiosis have showed that the benefits and costs of harboring symbionts in the social amoeba vary according to environmental contexts such as soil conditions (37) and food availability (16, 38). However, these studies primarily focused on contemporary interactions rather than their historical foundations. It also seems likely that hosts infected with different bacteria will evolve differently, at least if the infections persist long enough. If such evolution does occur, the different histories may impose historical contingencies on the host such that they evolve differently even when the symbionts are removed, and their environments are the same.

Here, we investigated the *Dictyostelium-Paraburkholderia* system to test whether *D. discoideum* strains show differential adaptations based on the history of symbiotic associations even after the symbionts are removed. We compared three traits between experimentally evolved non-symbiotic and symbiotic strains of wild collected *D. discoideum.* For each of the three life history traits, our results from univariate mixed models showed that *D. discoideum* strains with no history of symbiosis did not show significant response to experimental evolution in any phenotypic traits (Table 1). Overall, strains with previous symbiotic association showed a statistically significant response to experimental evolution without their bacterial endosymbionts for each of the traits (Table 1), indicating that partnering with a *Paraburkholderia* symbiont may alter the evolutionary potential of *D. discoideum* compared to non-symbiotic strains. We observed symbiont-specific patterns on the host *D. discoideum* traits by taking into consideration which species of *Paraburkholderia* symbionts the host strains lost and evolved without (Supplementary Table S2). Strains that lost and evolved without *P. agricolaris* showed significant increase in cell proliferation (Figure 1A), a significant decrease in slug migration (Figure 1B) but no change in spore production (Figure 1C). *P. hayleyella* partners showed an increase in both cell proliferation (Figure 1A) and spore production (Figure 1C) but no change in slug migration (Figure 1B) when evolved in the absence of the symbiont. Interestingly, *D. discoideum* strains that lost and evolved without *P. bonniea* only showed a significant decrease in the slug migration distance (Figure 1B), while cell proliferation and spore production were not affected (Figure 1A and Figure 1C).

The univariate mixed-effects models and standardized effect-size comparisons indicate that evolutionary responses to experimental symbiosis were detectable but modest and context dependent. Once corrected for multiple comparisons, many effects were small in magnitude, consistent with weak selection or limited statistical power at the level of individual traits. Taken together, the univariate and standardized analyses reveal the outlines of evolutionary change but only part of its structure.

The limited significance of single-trait responses, despite consistent effect-size patterns, suggests that evolutionary shifts may be subtle yet coordinated across traits. This possibility motivated a multivariate approach, in which correlated trait responses can jointly express an underlying axis of evolutionary divergence that is not apparent when traits are considered independently. The multivariate analyses reinforce this interpretation by demonstrating that evolved and ancestral lines differ significantly when all traits are considered jointly. Results from multivariate analyses revealed that symbiont history not only influenced the magnitude of evolutionary responses but also their directionality in multivariate trait space. The structure of trait variation revealed by our PCA suggests that symbiont history influences the evolutionary trajectory of *D. discoideum* across multiple functional axes. The primary axis of variation (PC1) has same-sign loadings for all three fitness components, and likely represents overall response to experimental evolution and separates ancestral strains from their descendants. The second axis (PC2) highlights a potential life-history trade-off between reproduction (sporulation) and dispersal (migration). To quantify the magnitude of evolutionary response that the loss of different symbiont species may have resulted in their hosts, we estimated evolutionary trajectories as vectors between the centroids of ancestral and evolved lineages on the PCA for the strains (Figure 2A). We further estimated angular deviations between the vectors to quantify the degree by which the evolutionary trajectories varied across strains with different history of carrying symbionts in the wild (Figure 2B). Strains that were previously hosts of *P. agricolaris* evolved in a direction similar to non-symbiotic strains but showed twice as much response to experimental evolution (Figure 2C). Strains that lost and evolved without *P. hayleyella* and *P. bonniea* showed the largest magnitude of evolutionary response but in directions away from each other (Figure 2A-C). Additionally, *P. bonniea* host strains showed an evolutionary trajectory that deviated from all other strains (Figure 2B). Taken together with the univariate results, these results suggest that evolutionary adaptation to symbiosis (and loss thereof) proceeds through coordinated modification of multiple, functionally connected traits rather than through isolated changes in single trait optimality. Such multivariate evolution may reflect both developmental and physiological coupling among traits and the complex selective environment imposed by microbial partners.

In addition to generally showing historical contingency, our results showed specific insights into the *Dictyostelium-Paraburkholderia* system. All three species of *Paraburkholderia* are facultative endosymbionts of the social amoeba that can be easily gained and lost, at least in laboratory conditions. When released from their symbionts, it is possible that the hosts evolve back towards an optimal phenotype for uninfected strains. Loss of *P. agricolaris*, with its full-sized genome, resulted in evolutionary responses that were quite like the non-symbiotic strains whereas loss of either of the two genome-reduced species led to evolutionary trajectories that were very different from that of the non-symbiotic strains. Genome reduction is indicative of symbionts being more dependent on the host and more highly adapted to life inside the host. This might lead to infections being more persistent and causing more change during the course of an infection. *P. agricolaris*, being less adapted to life inside amoebas, may cause shorter infections and therefore cause relatively little amoeba evolution during an infection and after being lost.

Although the reduced genomes of *P. bonniea* and *P. hayleyella* strongly suggest that these species have a longer history as symbionts than does *P. agricolaris*, we do not know how long individual infections persist in nature. However, some insight into this question is provided our results. *D. discoideum* periodically undergoes sexual reproduction (39), so long-term adaptations to each of the symbionts should get shared across the population. It is therefore short-term adaptations, due to a strain’s recent experience with a particular symbiont, that are expected to cause different trajectories after symbiont loss. The fact that we get different trajectories suggests that the most recent infection (the one we detected in each collected amoeba strain) persisted long enough for evolution to change amoeba physiologies in path-dependent ways.

The evolution of obligate symbiosis in a variety of taxa illustrates historical contingencies due to presence or absence of partners. It has been described as an evolutionary addiction or a rabbit hole of dependence (39,40). Loss of an obligate partner changes evolutionary trajectories – it leads to the inability to evolve. Our study extends this phenomenon to less dependent, facultative symbionts. We showed that a history of different bacterial partners can influence the evolution of distinct combinations of host traits, producing historical contingencies that can be symbiont specific. Even though the partners can separate, our study suggests that historical symbiotic relationships could create lasting effects on host evolutionary trajectories, long after those relationships are disrupted.

## Methods

### Study organism and Dataset

The dataset used for this study comes from an experimental evolution study (Larsen et al 2023) where *D. discoideum* wild-collected ancestral strains were experimentally evolved in the laboratory. The original experiment focused on understanding whether cheating in the social amoebae atrophies when selection for cooperation is reduced. Before experimental evolution, all strains were treated with antibiotics to remove bacterial endosymbionts. Therefore, *D. discoideum* strains that were collected with *Paraburkholderia* spp. in the wild (24) evolved without their bacterial partners during laboratory culture for ∼200–300 cellular generations. In a separate study, we made mixes between and among the strains and measured phenotypic traits to test whether cell lineages evolved in isolation from each other would generate incompatibilities due to a failure in coordination (31). Here we test whether a history of symbiosis explains the trajectory of phenotypic trait evolution in these experimental lineages. The dataset is available in the open-access github repository: https://github.com/jahanisrat/SymbiontLoss

We divided the *D. discoideum* strains into four groups based on their symbiont histories: (1) non-symbiotic strains that lacked symbionts when collected, (2) strains cured of *Paraburkholderia agricolaris*, (3) strains cured of *Paraburkholderia bonniea*, and (4) strains cured of *Paraburkholderia hayleyella*. We included three trait variables in this study that represent various aspects of performance or fitness throughout the *D. discoideum* life cycle. Briefly, the measurements from the previous study are described below:

1. Cell Proliferation: Measurements for this trait represent values from cell count assay. Total cell numbers were calculated after inoculating spores on SM/5 plates. Cells were counted prior to aggregation to isolate effects on vegetative growth from potential differences in sporulation efficiency. This assay specifically quantified proliferation capacity during the unicellular phase.
2. Slug Migration: Distances traveled by slugs towards a directional light source were measured in the slug migration assay. Spores (10^7^) were inoculated, with bacterial food, along one end of nutrient-free plates and given 8 days to migrate toward a pinhole light source on the opposite end. Migration distance was measured in centimeters from the starting line.
3. Spore production: Measurements for this trait represented spore counts on the non-nutrient migration plates for the same number of starting spores. Total spores produced by fruiting bodies on the plate were quantified by counting spores after migration. This measure captures the ability of spores to undergo the multicellular aggregate cycle, and complete successful culmination of fruiting bodies.

## Statistical Analysis

We used R (version 4.4) to perform statistical analyses. Data visualizations were created using the ggplot2 package.

For each of the univariate traits (cell proliferation, slug migration, and spore production), we fitted linear mixed effect models using the package *lme4* (40) and robust mixed effect models using *robustlmm* (41). We generated three models:

M0) Mixed model without a effect for the history of symbiosis

Trait ∼ treatment + (1|Strain/Line)

M1) Mixed model used to compare symbiotic vs non-symbiotic strains

Trait ∼ treatment * symbiosis + (1|Strain/Line)

where Trait was either Cell Proliferation, Slug Migration or Spore Production

M2) Mixed model formula used to compare strains with distinct symbionts

Trait ∼ treatment * symbiont + (1|Strain/Line)

where ‘Trait’ was either Cell Proliferation, Slug Migration or Spore Production. The predictor ‘treatment’ corresponds to the categorical variable for whether the experimental line is Ancestral or Evolved in the laboratory. ‘symbiosis’ corresponds to the categorical variable for whether the strain has a history of carrying symbionts (Yes) or is a non-symbiotic strain (No); ‘symbiont’ corresponds to the species of symbiont lost (None, *P. agricolaris*, *P. bonniea*, *P. hayleyella*)

We compared model performance indices among the three models above using the package *performance* (*42*) and summarized the results in Table S1. We estimated marginal means from the above models for post-hoc hypothesis testing using the package *emmeans* (43). Table 1 in main text summarizes the pairwise contrasts for Model 1. Table S2 in supplementary material summarizes the pairwise contrasts for Model 2. p-values from Table S2 were used for significance stars in Figure 1A, 1B and 1C in main text.

To determine how traits evolved in response to symbiont loss, we calculated a standardized measure of phenotypic change (Figure 1D in main text). We estimated standardized effect size using Cohen’s d, which allows us to make comparisons across traits easier (44–46). Cohen’s d provides a way to compare the effect size of differences between populations in a standardized manner. For example, it shows whether changes in traits like spore production or cell proliferation are small, moderate, or large.

We measured Cohen’s d for strains with the same symbiont history using the following formula:

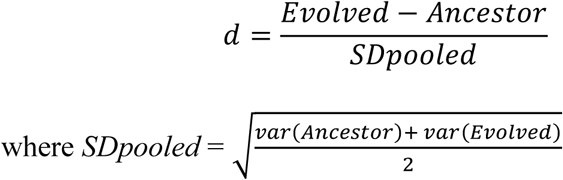

We used Multivariate Analysis of Variance (MANOVA) to test whether evolutionary responses differed among strains with distinct symbiont histories while accounting for correlations between traits. The model included three response variables (cell proliferation, slug migration, spore numbers) with treatment (Ancestor/Evolved) and cured symbiont species (None, *P. agricolaris*, *P. hayleyella*, *P. bonniea*) as predictors. MANOVA tests the null hypothesis that group mean vectors are sampled from the same multivariate normal distribution. Significant interaction between treatment and symbiont history would indicate that evolutionary trajectories differed based on symbiont type.

We evaluated MANOVA results using three test statistics: (1) Wilks’ Lambda (λ): represents the proportion of variance unexplained by group differences, (2) Pillai’s Trace: sums the proportion of variance explained on each discriminant function, (3) Roy’s Largest Root: maximum eigenvalue of the effect matrix. We transformed these test statistics to approximate chi-square distributions for significance testing (Table S3). Higher chi-square values indicate stronger evidence for group differences in evolutionary response.

To test whether symbiont history influences evolutionary trajectories independent of strain genetic background, we performed Permutational Multivariate Analysis of Variance (PERMANOVA) on trait data using Euclidean distance. We performed the analysis both with and without stratification by Strain to test whether strain-specific effects affected overall evolutionary change patterns and model significance.

## Author statements

### Author contributions

I. J. conceived of the study. T. J. L. experimentally evolved the *D. discoideum* strains. I. J., T. J. L., J.E.S., and D.C.Q. contributed to the statistical analyses, writing, and editing the manuscript.

### Conflicts of interest

The author(s) declare that there are no conflicts of interest.

### Funding information

This material is based upon work supported by the National Science Foundation under grant numbers NSF DEB-1753743 and DEB-2237266. I. J. was additionally supported by a Ph.D. fellowship from the McDonnell International Scholars Academy, WUSTL.

### Ethical approval

No experimental work with humans or animals.

## Acknowledgements

We thank Trey. J. Scott for helpful comments on the statistical analyses. Insightful discussions with Calum J. Stephenson, P. M. Shreenidhi, and other members of the Queller-Strassmann Lab helped improve this manuscript.

## SUPPLEMENTARY TABLES

**Table S1.** Test statistics comparing whether the effect of evolution differs between symbiosis histories. Estimates report: custom interaction contrasts on *Dictyostelium discoideum* trait evolution: (Evolved−Ancestral) _Symbiotic_ − (Evolved−Ancestral) _Non-symbiotic_.

| Trait | Estimate | SE | z.ratio | p.value |
| --- | --- | --- | --- | --- |
| Cell proliferation | 0.045 | 0.0996 | 0.452 | 0.651 |
| Slug Migration | –0.544 | 0.605 | –0.900 | 0.368 |
| Spore production | 0.065 | 0.0932 | 0.692 | 0.489 |

**Table S2A.** Model summaries for mixed effect models for fixed effects of experimental evolution (Treatment), a history of symbiosis (Symbiosis), and their interaction (Treatment × Symbiosis) across three traits – cell proliferation, slug migration and spore production.

| Trait | Effect | Chisq | DF | Pr(>Chisq) |
| --- | --- | --- | --- | --- |
| Cell Proliferation | Treatment | 1.0287 | 1 | 0.310 |
|  | Symbiosis | 0.0004 | 1 | 0.984 |
|  | Treatment × Symbiosis | 0.1196 | 1 | 0.730 |
| Slug Migration | Treatment | 0.2461 | 1 | 0.620 |
|  | Symbiosis | 2.7187 | 1 | 0.100 |
|  | Treatment × Symbiosis | 0.3480 | 1 | 0.555 |
| Spore Migration | Treatment | 0.1271 | 1 | 0.721 |
|  | Symbiosis | 0.0241 | 1 | 0.876 |
|  | Treatment × Symbiosis | 0.6582 | 1 | 0.417 |

**Table S2B.** Model summaries for mixed effect models for fixed effects of experimental evolution (Treatment), a history of carrying a particular symbiont (Symbiont), and their interaction (Treatment × Symbiont) across three traits – cell proliferation, slug migration and spore production.

| Trait | Effect | Chisq | DF | Pr(>Chisq) |
| --- | --- | --- | --- | --- |
| Cell Proliferation | Treatment | 1.288 | 1 | 0.256 |
|  | Symbiont | 3.284 | 3 | 0.349 |
| | Treatment × Symbiont | 9.232 | 3 | 0.026 |
| Slug Migration | Treatment | 0.268 | 1 | 0.604 |
|  | Symbiont | 3.539 | 3 | 0.315 |
| | Treatment × Symbiont | 4.914 | 3 | 0.178 |
| Spore Migration | Treatment | 0.1247 | 1 | 0.724 |
|  | Symbiont | 6.992 | 3 | 0.072 |
| | Treatment × Symbiont | 2.118 | 3 | 0.548 |

**Table S3.**
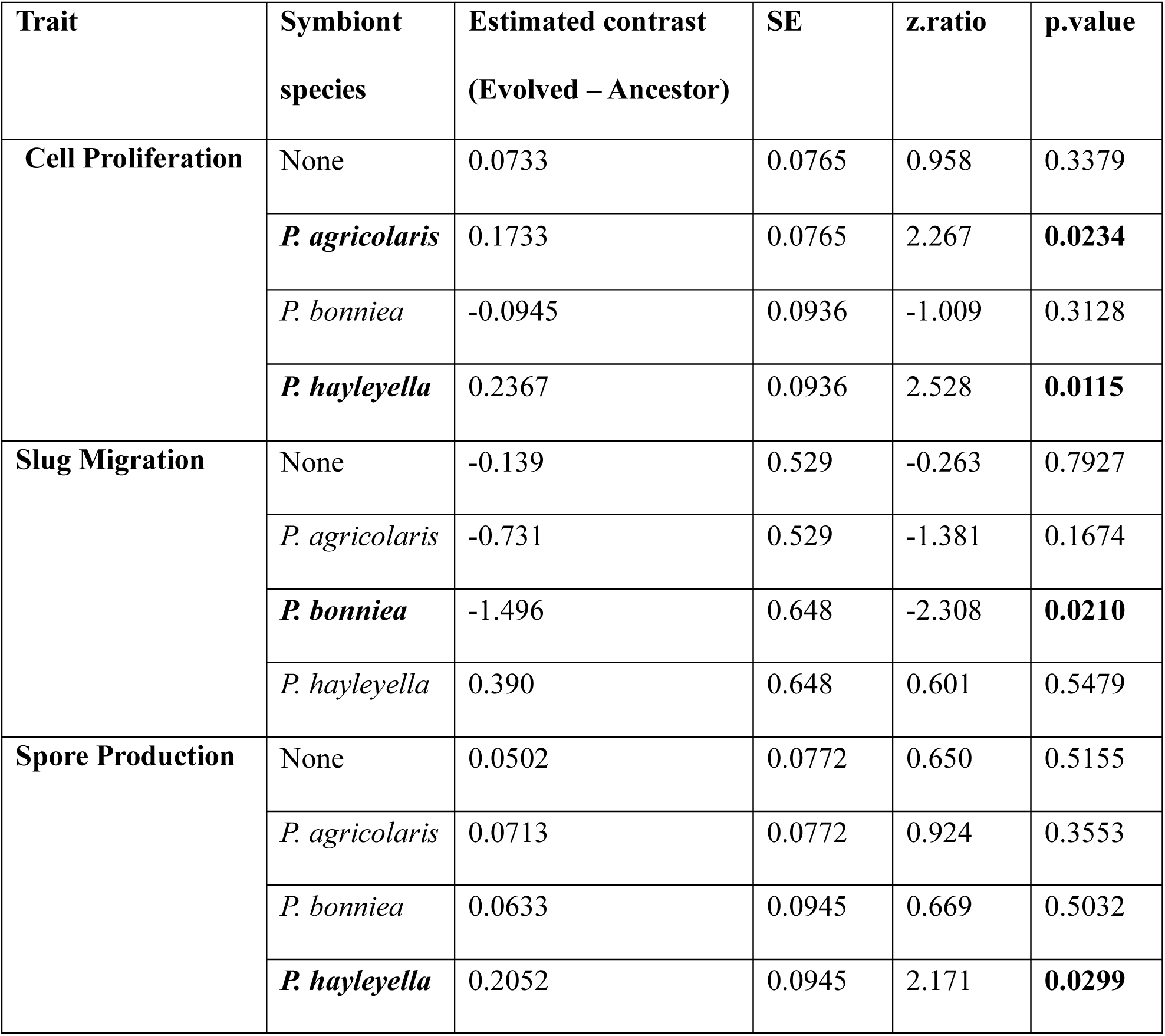
Statistics from pairwise contrasts between Ancestors and Evolved Lineages for *Dictyostelium discoideum* strains that carried distinct species of symbionts. The traits cell proliferation and spore production were calculated as log transformed number of cells and spores respectively. Slug migration was measured as distance traveled towards light in centimeters. *p-values* are FDR-adjusted for six comparisons per trait.

**Table S4.** Pairwise interaction contrasts of the evolutionary effect across symbiont identities. Each estimate represents the difference between two symbionts in the size of the evolutionary change (Evolved − Ancestor) for the specified trait. *p-values* are FDR-adjusted for six comparisons per trait.

| Trait | Treatment contrast | Symbiont pair | Estimate | SE | z.ratio | p.value |
| --- | --- | --- | --- | --- | --- | --- |
| Cell proliferation | Evolved – Ancestor | None – *P. agricolaris* | –0.100 | 0.108 | –0.93 | 0.426 |
|  |  | None – *P. bonniea* | 0.168 | 0.121 | 1.39 | 0.265 |
|  |  | None – *P. hayleyella* | –0.163 | 0.121 | –1.35 | 0.265 |
|  |  | *P. agricolaris* – *P. bonniea* | 0.268 | 0.121 | 2.22 | 0.080 |
|  |  | *P. agricolaris* – *P. hayleyella* | –0.063 | 0.121 | –0.52 | 0.600 |
|  |  | *P. bonniea* – *P. hayleyella* | –0.331 | 0.132 | –2.50 | 0.074 |
| Slug migration | Evolved – Ancestor | None – *P. agricolaris* | 0.592 | 0.749 | 0.79 | 0.515 |
|  |  | None – *P. bonniea* | 1.357 | 0.837 | 1.62 | 0.315 |
|  |  | None – *P. hayleyella* | –0.529 | 0.837 | –0.63 | 0.528 |
|  |  | *P. agricolaris* –<br>*P. bonniea* | 0.765 | 0.837 | 0.92 | 0.515 |
|  |  | *P. agricolaris* –<br>*P. hayleyella* | –1.120 | 0.837 | –1.34 | 0.361 |
|  |  | *P. bonniea* – *P.*<br>*hayleyella* | –1.886 | 0.917 | –2.06 | 0.238 |
| **Spore**<br>**production** | Evolved – Ancestor | None – *P.*<br>*agricolaris* | –0.021 | 0.109 | –0.19 | 0.947 |
|  |  | None – *P.*<br>*bonniea* | –0.013 | 0.122 | –0.11 | 0.947 |
|  |  | None – *P.*<br>*hayleyella* | –0.155 | 0.122 | –1.27 | 0.577 |
|  |  | *P. agricolaris* –<br>*P. bonniea* | 0.008 | 0.122 | 0.07 | 0.947 |
|  |  | *P. agricolaris* –<br>*P. hayleyella* | –0.134 | 0.122 | –1.10 | 0.577 |
|  |  | *P. bonniea* – *P.*<br>*hayleyella* | –0.142 | 0.134 | –1.06 | 0.577 |

**Table S5.** Chi-square approximations for multivariate test statistics assessing the effects of treatment, symbiont history, and their interaction on *Dictyostelium discoideum* trait evolution.

| Variable | Test Statistic | Value | Chi-Square Approx. | p-value |
| --- | --- | --- | --- | --- |
| **Treatment** | Wilks' Lambda | 0.82 | 22.47 | <0.001 |
| **Treatment** | Pillai's Trace | 0.18 | 61.11 | <0.001 |
| **Treatment** | Roy's Largest Root | 0.22 | 25.51 | 0.002 |
| **Symbiont** | Wilks' Lambda | 0.43 | 96.30 | <0.001 |
| **Symbiont** | Pillai's Trace | 0.66 | 226.48 | <0.001 |
| **Symbiont** | Roy's Largest Root | 0.90 | 104.25 | <0.001 |
| **Treatment × Symbiont** | Wilks' Lambda | 0.83 | 20.62 | 0.014 |
| **Treatment × Symbiont** | Pillai's Trace | 0.17 | 58.23 | <0.001 |
| **Treatment × Symbiont** | Roy's Largest Root | 0.16 | 18.53 | 0.029 |

Note: Chi-square approximations were calculated as: $-(n - 1 - (p + df)/2) * \log(\text{Wilks' Lambda})$ for Wilks' Lambda, $\text{Pillai's Trace} * \min(p, df) * (n - 1 - (p + df)/2)$ for Pillai's Trace, and $\text{Roy's Largest Root} * (n - p - 1)$ for Roy's Largest Root, where n is the sample size, p is the number of response variables, and df is the degrees of freedom.

## Notes

https://github.com/jahanisrat/SymbiontLoss

### Competing Interest Statement

The authors have declared no competing interest.

